# Wnt activation in mature dermal adipocytes leads to lipodystrophy and skin fibrosis via ATGL-dependent lipolysis

**DOI:** 10.1101/2024.11.16.623972

**Authors:** Qiannan Ma, Ella X. Segal, Claire Reynolds, Suneeti R. Madhavan, Rachel H. Wyetzner, Megan Gregory, Radhika P. Atit

## Abstract

**Objective:** Accumulation of extracellular matrix (ECM) and dermal adipocyte lipodystrophy occurs during skin fibrosis, which compromises the skin’s flexibility and function. We recently showed that sustained Wnt activation in dermal progenitor cells leads to fibrotic ECM thickening in the dermis and lipodystrophy of dermal white adipose tissue (DWAT). The aim of this study was to test if Wnt/β-catenin signaling in mature dermal adipocytes directly leads to lipodystrophy and impact skin fibrosis.

**Methods:** We developed a genetically lineage traceable, DWAT specific, inducible and reversible mouse model of Wnt activation (Adipo-β-cat^istab^) in the dorsal skin. We analyzed the DWAT lipid droplet size, cell identity, and affect on ECM accumulation and remodeling in skin fibrosis. Adipocyte triglyceride lipase (ATGL) is the key rate limiting enzyme of the lipolysis pathway, which is a biological process of breaking down triglyceride stores in adipocytes. The *Atgl* gene was conditionally deleted in mature dermal adipocytes to test the requirement of the lipolysis pathway in the Wnt-induced lipodystrophy (Adipo-β-cat^istab^; *Atgl^fl/fl^*).

**Results:** Here, we utilize mouse genetic models with lineage tracing to show that Wnt activation in mature dermal adipocytes is sufficient to induce adipocyte lipodystrophy and fibrotic collagen remodeling. Upon withdrawal of adipocyte-restricted Wnt activation, lipodystrophy and fibrosis were reversed. Mechanistically, we find that Wnt activation stimulates Adipose Triglyceride Lipase (ATGL)-mediated lipolysis pathway. We found *Atgl* in dermal adipocytes is functionally required for Wnt-induced lipodystrophy in the DWAT and fibrotic remodeling.

**Conclusion:** Collectively, this study demonstrates that Wnt activation in dermal adipocytes promotes lipolysis and may be a novel therapeutic avenue for preventing and reversing lipodystrophy and skin fibrosis.

## Introduction

In chronic tissue fibrosis, extracellular matrix proteins such as collagens accumulate, leading to increase in tissue stiffness and a-cellularity that can cause organ failure and morbidity (1). Fibrosis can occur in all most tissues including skin, lung, liver, heart, intestine and kidney (1,2). Despite its devastating impact on nearly 1 in 4 people and annualized incidence of major fibrosis related conditions in 1 in 20 people globally, no effective treatment exists to reverse fibrosis (2). During fibrosis, several tissues such as skin, liver, and lung have loss of lipid-filled cells or lipid depletion of adipocytes leading to lipodystrophy (3–7). However, the signals and mechanisms underlying fibrotic lipodystrophy are not fully understood.

In the skin, the dermal adipocytes form a distinct dermal white adipose tissue (DWAT) layer that sits beneath the ECM-rich dermis, making the skin an ideal system to understand the physiological mechanisms underlying lipodystrophy. Dermal fibroblasts are the key producers of ECM which provides structural integrity to the skin (8). DWAT contributes to thermoregulation, produces antimicrobial peptides to modulate immune response upon injury (9,10). The dynamic change in size of DWAT in homeostatic conditions occurs during adaptive thermogenesis, the hair follicle cycle, and wound healing (7,9,11). Skin fibrosis is marked by the buildup of extracellular matrix (ECM) due to heightened activation and proliferation of dermal fibroblasts, along lipodystrophy of the DWAT. These changes collectively lead to tissue stiffening and impaired function. Chronic skin fibrosis occurs in diseases such as systemic sclerosis (SSc), atopic dermatitis, diabetes, and psoriasis affecting 100 million people globally each year (12). DWAT lipodystrophy is a hallmark of acute fibrosis in wound healing and chronic fibrosis in human systemic scleroderma skin (SSc), but the impact of DWAT lipodystrophy on dermal fibrosis is unclear (13). To identify new strategies for preventing and reversing established fibrosis, we need to define the inducing signals that cause fibrotic lipodystrophy.

Of the many profibrotic pathways, the canonical Wnt/β-catenin signaling pathway stands out because it is a conserved profibrotic pathway across tissues and anti-adipogenic (14,15). Wnt/β-catenin signaling can upregulate profibrotic genes such as connective tissue growth factor (CTGF), TGF-β signaling components, smooth muscle actin (ACTA2), and matrix genes such as COL1A1 to cause ECM accumulation (16–19). In contrast, Wnt/β-catenin signaling activation leads to epigenetic silencing of *Ppar-g* and *Cebp-a* which are the central regulators of adipocyte fate (20,21). Previously, our lab showed Wnt/β-catenin signaling activation in Engrailed1^+^ fibro-adipocyte progenitors and their derivatives can cause ECM accumulation in the dermis and lipid depletion of the DWAT (18). Lipid homeostasis in adipocytes is a balance of lipid accumulation and lipid breakdown. Lipolysis is the physiological pathway that adipocytes use to breakdown triacylglycerol (TAG) stored in the lipid droplet by a sequence of lipases starting with Adipose Triglyceride Lipase (ATGL) which results in the release of fatty acid and glycerol (22,23). Emerging evidence suggests that factors or fatty acids from adipocytes can contribute to the fibrotic transformation and tissue dysfunction (2). Whether Wnt/β-catenin signaling is directly involved in adipocyte lipodystrophy and its impact on skin fibrosis is not known.

To determine if dermal adipocyte restricted Wnt activation is sufficient to cause lipodystrophy or skin fibrosis, we constructed an inducible and reversible genetic mouse model to activate Wnt signaling pathway in *Adiponectin*+ mature dermal adipocytes with indelible genetic lineage tracing. We found Wnt activation in mature dermal adipocytes was sufficient to cause dermal ECM remodeling, elevated proliferation in the skin, and lipodystrophy of DWAT without perturbing cell survival. These phenotypic changes require the ATGL-dependent lipolysis pathway and can also be rescued upon withdrawal from Wnt activation. Collectively, these data show that skin fibrosis associated lipodystrophy and ATGL-dependent lipolysis may be a new therapeutic target in fibrosis treatment.

## Materials and Methods

*AdiponectinCre*ER (24) (Jax stock 024671); Rosa26mTmG (25) (Jax stock 007676 respectively); Rosa26rtTA-EGFP (26)(Jax Stock 005572); TetO-deltaN89*β*-catenin (27); *Atg*l^flox/flox^ (28) (Jax stock 024278) mice were used for generating the dermal-adipocyte restricted, inducible and reversible Wnt signaling activation and conditional deletion of lipolysis mouse model. Case Western Reserve Institutional Animal Care and Use Committee approved all animal procedures in accordance with American Veterinary Medical Association guidelines protocol 2013-0156, approved 21 Dec. 2021, Animal Assurance No. A3145-01 at Case Western. Mice are maintained and bred on a mixed genetic background in the Animal Resource Center (ARC) at Case Western Reserve University. Males and females are included in all experiments. Phenotypic analysis was based on at least two to five litters of mice with litter-matched mutants and controls.

To spatially and temporally restrict activation of Wnt signaling pathway in mature *Adiponectin*-CreER dermal adipocytes in mouse dorsal skin, 100μL of 5mg/mL tamoxifen in 100% ethanol (ApexBio B5965) was applied topically to shaved dorsal skin of weaned mice at 21-day (P21) and 22-days old (P22) (7,29). *Adiponectin*-CreER line was used to recombine both Rosa26 membrane-targeted tandem dimer Tomato/membrane-targeted green fluorescent protein (R26mT/mG) and Rosa26 reverse tetracycline regulator transactivator (R26rtTA). For induction of β-catenin-myc tagged transgene expression, the key transducer of canonical Wnt signaling pathway, in the *Adiponectin-* CreER/+; R26rtTA/+;R26mTmG;TetO-deltaN89*β*-catenin/+ mice (Adipo-β-cat^istab^), 23-day old (P23) mice were given 6g/kg doxycycline chow (Envigo-Harlan) and 2 mg/mL doxycycline (Fisher Scientific 446061000) water daily for 5 (P23-28) and 10 days (P23-33). In reversal experiments after Wnt activation regimen, mice were switched to 10 days of regular chow and water (P34-P44). To preferentially and genetically inhibit lipolysis in mature dermal adipocytes on mouse dorsal skin, *Adiponectin-*CreER line was used to delete *Atgl^fl/fl^*as previously shown (7,30–32)*. Atgl^fl/fl^* mice were bred into the Adipo-β-cat^istab^ background and the Wnt activation regimen was performed with doxycline in chow and water to Adipo-β-cat^istab^; *Atgl^fl/fl^* mice for 5 or 10 days and skin was analyzed at specific time points at P28, P33. Control groups were CreER negative or bigenic (Fig.1,4) with similar treatments as the experimental group.

**Figure 1.**
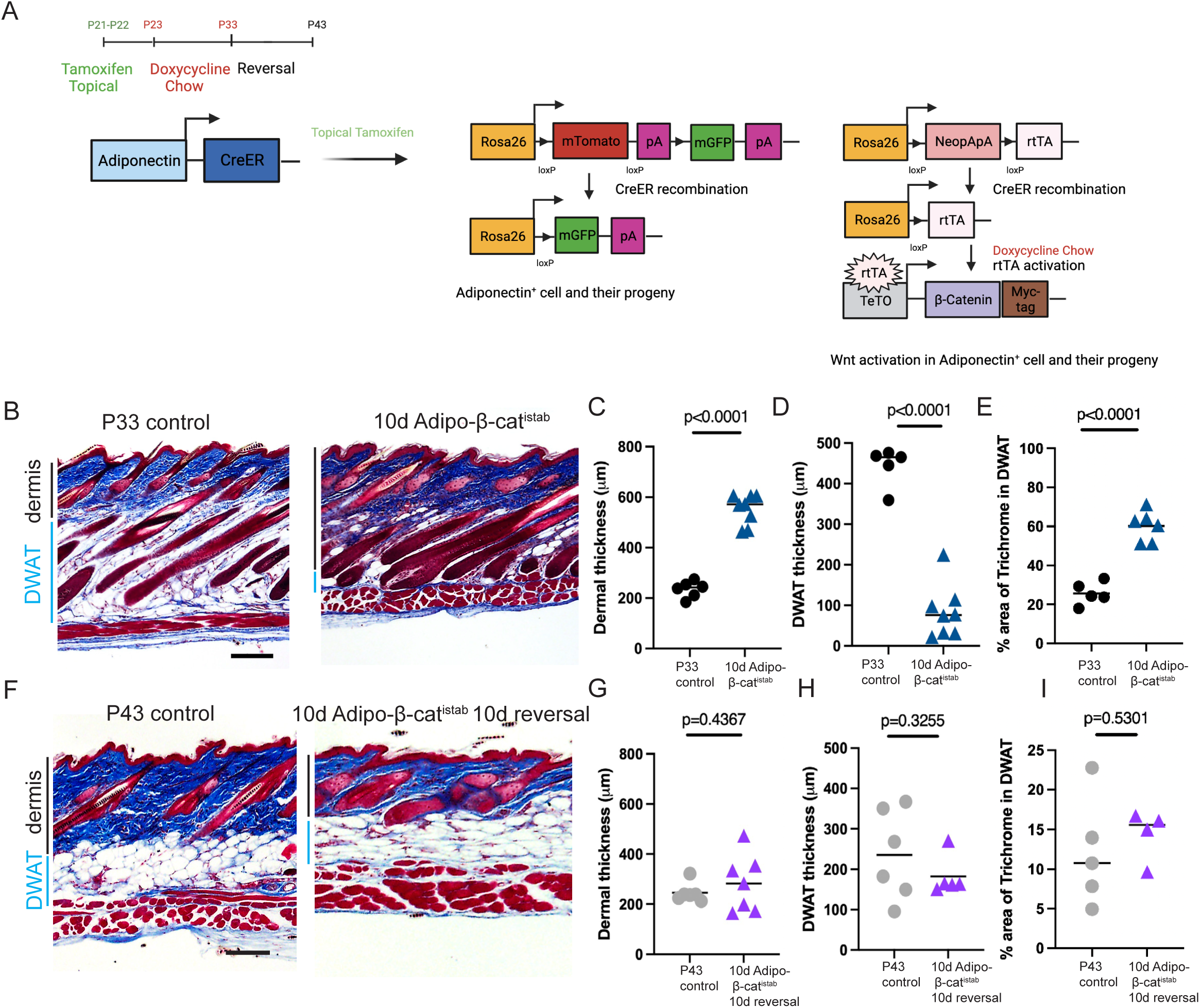
Adipo-β-cat^istab^ leads to inducible and reversible dermal thickness increase and DWAT thickness decrease. **A.** Experimental design: 21-d-old (P21) control and Adipo-β-cat^istab^ mice were treated with topical tamoxifen on dorsal skin and fed doxycycline containing chow and water for inducible-reversible Wnt activation in mature DWAT cells. Skin was harvested after 10 days (P33) or subsequently after 10 days of withdrawal from Wnt activation (P43). **B.** Masson’s trichrome staining of mouse dorsal skin cut in sagittal plane from (P33) control, 10 days Adipo-β-cat^istab^. (n=6-8). **C., D.** Quantification of mouse dorsal dermal thickness and DWAT thickness in (**B**) (n=5-8). **E**. Quantification of area percentage of trichrome in Masson’s Trichrome staining of DWAT for (P33) control and 10 days Adipo-β-cat^istab^. (n=5-6) **F**. Masson’s trichrome staining of mouse dorsal skin for (P43) control and 10 days Adipo-β-cat^istab^; 10 days reversal. **G-H.** Quantification of mouse dorsal dermal thickness (n=6-7) and DWAT thickness (n=5-6) in (**F**). **I.** Quantification of area percentage of blue in Masson’s Trichrome staining of DWAT for (P43) (n=5-6). Scale Bar=200μm, black and blue bars are the dermal (n=5-8) and DWAT thickness. *P* values were calculated with unpaired, two-tailed *t*-test with Welch’s correction and a *P* value of <0.05 is considered significant.

### Histological, Immunostaining and morphometrics

Samples were fixed in 10% formalin followed by paraffin embedding for Masson’s trichrome, Picrosirius Red (Electron Microscopy Sciences), Biotin-Collagen Hybridizing peptide (B-CHP) (BIO300, 2μM, 3Helix Company) as per manufacturer or previously described (18,33).

Immunohistochemistry staining with Myc-tag (ab9106, 1:500; Abcam), phospho-HSL (Ser 565) (4137, 1:1600; Cell Signaling) primary antibodies was performed as previously described except tertiary amplification was done with NeutrAvidin, (Vector Laboratories) (18). Bright field images were taken by an Olympus BX60 microscope with a digital camera (DP70, Olympus). Masson’s trichrome staining images were taken at 4x objective (Olympus UPIanFI 4x/0.13) by Cell Sens Entry software (Version 1.5, Olympus Corporation 2011). Flash frozen fresh skin was folded with the DWAT side inside and samples were embedded in O.C.T and cut sagittally at 20 microns. Immunofluorescence staining on frozen sections after 4% PFA fixation was performed with primary antibodies GFP (ab13970, 1:1000; Abcam), Perilipin1(PLN1) (ab3526, 1:500; Abcam), Ki67 (ab15580, 1:500; Abcam) and with species-specific Alexa-fluor secondary antibodies (Invitrogen) as described in detail in Supplementary methods. Programmed cell death was visualized with TUNEL assay according to manufacturer’s instructions for TUNEL TMR red kit (11767291910, 5% TUNEL enzyme; Roche Diagnostic Corp).

Fiji/ImageJ (National Institute of Health, Bethesda, MD) was used in phenotypic data quantification analysis for dermal thickness, dermal white adipose tissue thickness, cell counting, collagen amount analysis and individual adipocyte area measurements. Quantification of skin compartment thickness represents the average of three different locations in 3-6 sections/mouse. Pipelines and quantification of cells expressing specific markers is described in supplementary methods. The ROIs from polarized PSR images analyzed through TWOMBLI in Fiji/ImageJ (34). Line masks of the matrix network were generated by the Fiji Ridge Detection tool. According to the line mask, these ECM metrics were calculated from six ROIs, three different sections in three different regions of the treated skin per animal: curvature, fractal dimension, number of branch points, total length, number of endpoints, lacunarity, and alignment. The ROIs were converted into 16-bit grayscale by Fiji and input into Alignment by Fourier Transform (AFT) (35) by MATLAB (Mathworks, v2023b) for alignment analysis. A full description of the materials and methods is provided in the online supplementary materials.

## Results

### Wnt signaling activation in mature dermal adipocyte is sufficient to induce fibrotic remodeling in the whole skin

To determine if Wnt activation in the mature dermal adipocytes in the mouse dorsal skin is sufficient to induce skin fibrosis, we generated a new mouse model with lineage-tracing combined with an inducible and reversible expression of stabilized β-catenin-Myc tag (Adipo-β-cat^istab^) (**Fig.1A**) (7,24,26,27). Lineage tracing of mature dermal adipocytes by topical tamoxifen inducible *AdiponectinCreER; R26mT/mG* reporter revealed that 70% of adipocytes in the DWAT layer were GFP positive in the dorsal skin and 20%-30% GFP positive adipocytes in the other fat depot, which is consistent with previous reports (**Fig. 2B, Supplementary Fig. S1A, B**) (7). We did not observe GFP expression in the dermis (**Supplementary Fig. S1C**). After 5 days of Wnt activation in the Adipo-β-cat^istab^, we found nuclear expression of β-catenin-Myc-tag in at least 20-65% of adipocytes in a fixed field in the DWAT layer (**Supplementary Fig. S2A-C**). qPCR analysis of DWAT tissue showed a three-fold elevated expression of Axin2 mRNA, a transcriptional target of the canonical Wnt signaling pathway. (**Supplementary Fig. S2D**). Together these data indicate that *Adiponectin*-CreER/+ can efficiently activate Wnt signaling in DWAT of tamoxifen-treated Adipo-β-cat^istab^ mice.

**Figure 2.**
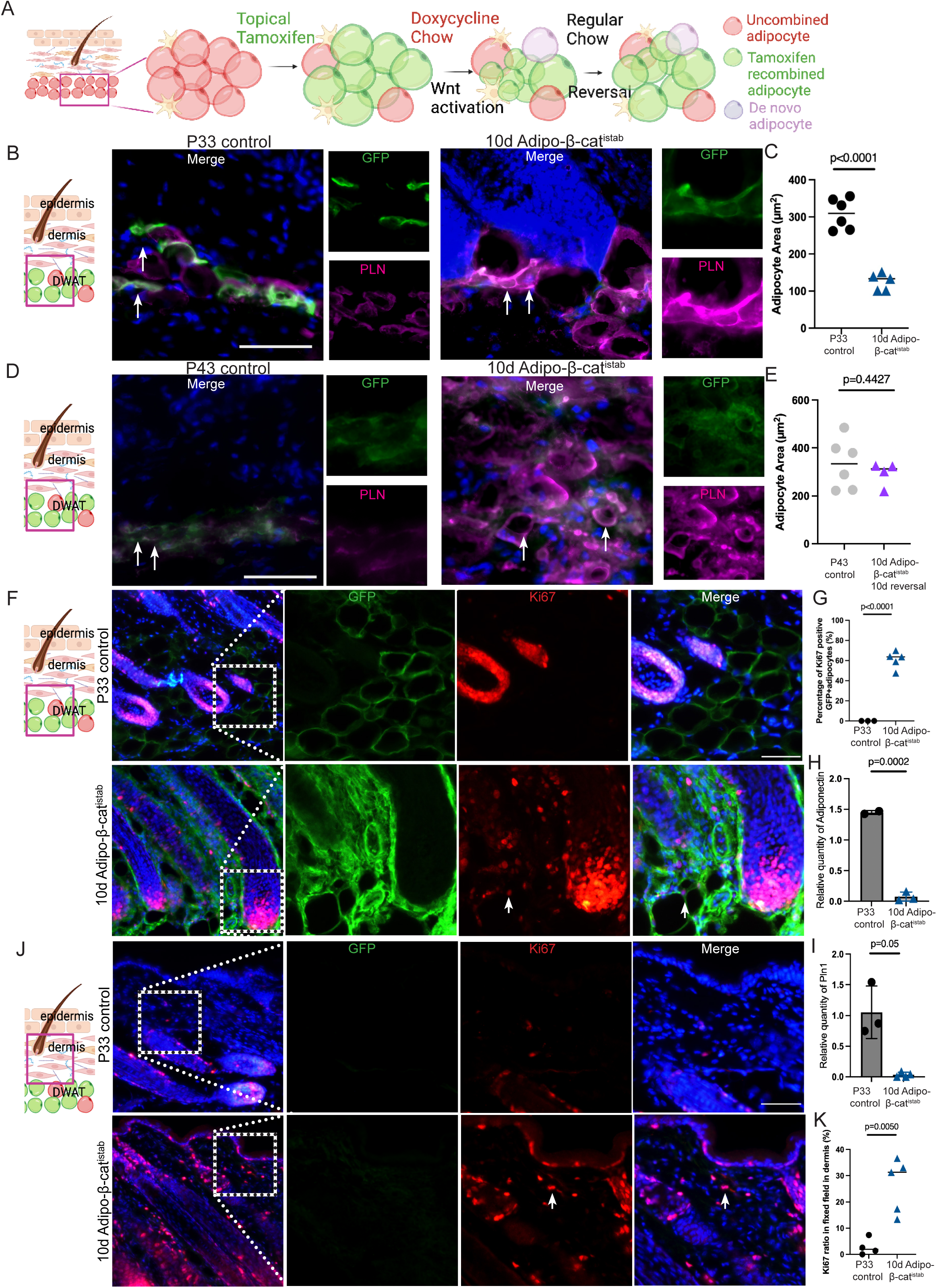
Adipo-β-cat^istab^ leads to inducible and reversible decreasing of mature dermal adipocyte size and whole skin cell proliferation response. **A.** Schematic for tamoxifen induced R26mTmG recombined mature adipocyte lineage labeling during Wnt activation and after subsequent withdrawal from Wnt activation. **B.**, **C.** GFP and Perilipin 1 immunofluorescence staining and quantification of PLN1+ vesicles in control (P33) and GFP+ vesicles in 10 days Adipo-β-cat^istab^ with low magnification on the left (n= 5). **D., E.** GFP and Perilipin 1 immunofluorescence and quantification of area of PLN1^+^ adipocytes in control (P43) and GFP+ vesicles in 10 days Adipo-β-cat^istab^; 10 days reversal (n=4-6). **F., G.** Ki67 and GFP immunostaining on P33 control and 10 days Adipo-β-cat^istab^ in the DWAT and quantification Ki67+ in GFP+ adipocytes per fixed field (n=3-5). **H.-I**. qPCR quantification of Perilipin 1 and Adiponectin on whole skin (n=2-4). **J., K.** Images with insets of Ki67 and GFP immunostaining in the skin and quantification of percent of Ki67^+^ cells in a fixed field dermis (n=4-5). DAPI=blue, GFP=green, Ki67=red, PLN1=purple, DAPI=blue. Scale bar= 100μm. White arrow stands for Ki67+ cell. *P* values were calculated with unpaired, two-tailed *t*-test with Welch’s correction and a *P* value of <0.05 is considered significant.

By 10 days of Wnt activation in mature dermal adipocytes with dietary doxycycline, we observed the earliest significant decrease of DWAT thickness, as well as increase in collagen amount in DWAT, accompanied by dermal ECM expansion (**Fig. 1B-E**). Although Wnt activation occurred in a subset of mature dermal adipocytes of Adipo-β-cat^istab^ mice, we still found a tissue level decrease in DWAT thickness and increase in dermal thickness. Next, we investigated if reversal of Adipo-β-cat^istab^ phenotypes occur upon subsequent withdrawal from Wnt activation. After 10 days of withdrawal from Wnt activation we observed a rescue of ECM expansion in the dermis and lipodystrophy as well as the collagen amount abundance in the DWAT in Adipo-β-cat^istab^ mice (**Fig. 1F-I).** Thus, our genetic inducible and reversible model demonstrates that sustained activation of the Wnt signaling pathway in the mature dermal adipocytes of the dorsal skin contributes to the two main characteristics of skin fibrosis: ECM expansion of dermis and lipodystrophy of DWAT.

### Wnt signaling activation in the mature dermal adipocytes leads to decrease in adipocyte size and identity with increase cell proliferation in the skin

To further investigate the effect of Wnt signaling activation on the mature dermal adipocytes when tissue level fibrotic changes occur, we measured the size of the *Adiponectin-*CreER*; R26mT/mG* lineage traced GFP^+^/PLIN1^+^ adipocytes after 10 days Wnt activation (**Fig. 2A**). Compared to the size of PLN1^+^ adipocytes in control mice, the size of the lineage-marked GFP^+^ dermal adipocytes were consistently decreased in Adipo-β-cat^istab^ mice and the adipocytes between hair follicles were flat in morphology and located adjacent to the hair shaft (**Fig. 2B, C, Supplementary Fig S1D)**. In the Adipo-β-cat^istab^ mice, size of non-recombined (GFP^_^) PLN1^+^ adipocytes appear larger than the lineage-marked GFP^+^ adipocytes, qualitatively. Subsequent withdrawal from Wnt activation in Adipo-β-cat^istab^ mice led to recovery and increase in size of lineage marked lipid depleted mGFP+ adipocytes (**Fig. 2D, E**). These data indicate that the size of the mature lipid filled dermal adipocytes is dynamic and this process is dependent on sustained Wnt activation.

Next, we tested the impact on cell survival after 5 and 10 days of Wnt activation in mature dermal adipocytes in dorsal skin by Terminal deoxynucleotidyl transferase dUTP nick-end labeling (TUNEL) assay. Cell death marker by TUNEL assay in the DWAT region was comparable in controls and Adipo-β-cat^istab^ skin (**Supplementary Fig. S3A, B).** In homeostatic conditions, the proliferation index of cells in the dermis of adult skin is very low and mature dermal adipocytes are terminally differentiated post-mitotic cells (36). We took advantage of lineage tracing to determine if mature dermal adipocytes dedifferentiated after Wnt activation induced lipodystrophy and re-enter the mitotic cycle (**Figure 2F-2K**). Compared to the morphology of mature lipid filled dermal adipocytes in the control skin, the DWAT layer in Adipo-β-cat^istab^ skin had mGFP+ dermal adipocyte cells that had shrunken and clustered around hair follicles and some are Ki67^+^(**Fig. 2F**). Next, we measured the proliferation index of cells in different parts of the skin by calculating the percent of Ki67+ cells in a fixed area (**Fig. 2F, Supplementary Fig. S3C, S3D**). As expected, the proliferation index was low in control skin dermis and DWAT regions (**Fig. 2F, 2G, 2J, 2K**). In contrast, Ki67^+^ nuclei were visible in the dermis and inside mGFP+ lineage marked cells in Adipo-β-cat^istab^ skin and quantification showed a significant increase of Ki67^+^ mGFP+ adipocyte number in the DWAT and Ki67+ cells in the dermal layer (**Fig. 2F, 2G, J, K**). We also observed a significant decrease in the mRNA expression of mature adipocyte markers, *Adiponectin* and *Perilipin1* in 10d Adipo-β-cat^istab^ mouse dorsal skin compared to control **(Fig 2H, 2I, Supplementary Fig. S3A and S3B**). Together, these data suggest that the Wnt activation in mature dermal adipocytes may lead to de-differentiation and reentry into the cell cycle and secondarily elevated cell proliferation in the dermis of the skin.

### Fibrotic stimulation of mature dermal adipocytes results in ECM remodeling and topographical changes in dermal collagen

Since we observed significant increase in area occupied by collagen in DWAT and increase in dermal thickness in Adipo-β-cat^istab^ mice (**Fig 1C, D**), we analyzed collagen remodeling by visualizing the location of unfolded collagen chains with Biotin Conjugate Collagen Hybridizing Peptide (B-CHP) (33). In control mouse skin, remodeling of collagen normally occurs in low levels in healthy dermis to maintain the collagen homeostasis and is nearly undetectable in the DWAT layer (**Fig. 3A**). We found that B-CHP significantly increased in the DWAT and dermis in the 10 days of Wnt activation in Adipo-β-cat^istab^ mice (**Figure 3B-3D**). Next, we investigated if collagen remodeling was reversible upon subsequent withdrawal from Wnt activation in Adipo-β-cat^istab^ mice. After 10 days of withdrawal from Wnt activation in mature dermal adipocytes, the B-CHP signal was rescued and comparable in both the DWAT and dermis in Adipo-β-cat^istab^ mice (**Fig. 3E-3H**). These results indicate that to the fibrotic remodeling of the ECM in the whole skin is dependent on sustained Wnt activation in the mature dermal adipocytes.

**Figure 3.**
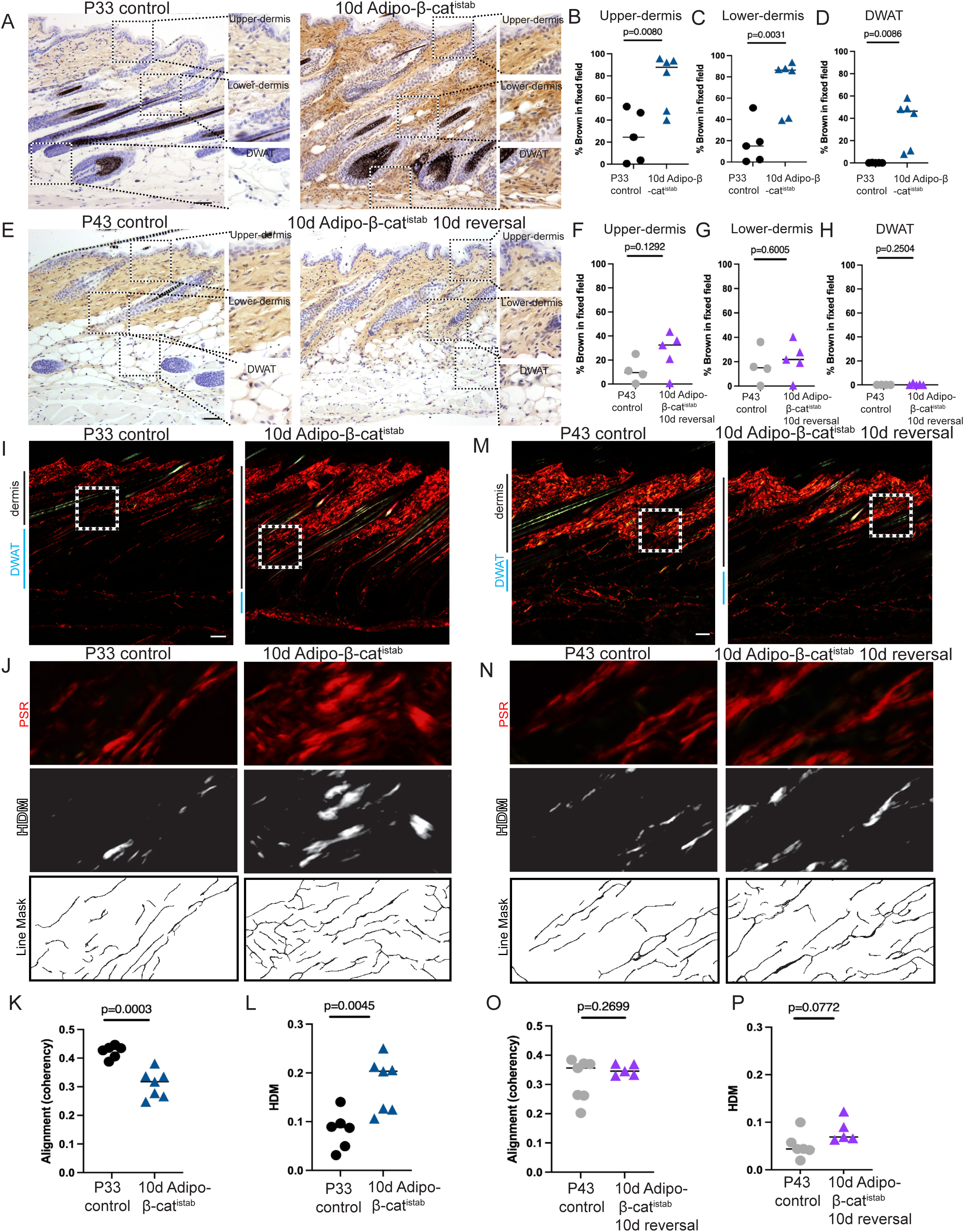
Adipo-β-cat^istab^ leads to inducible and reversible topographical changes and collagen remodeling in dermis. **A.** Collagen remodeling was visualized with collagen hybridizing peptide conjugated to biotin (B-CHP, brown) on (P33) control, 10 days Adipo-β-cat^istab^. **A.-D.** Representative views with insets of upper-dermis, lower-dermis and DWAT quantification of percent area B-CHP=brown in a fixed field in upper-dermis, lower dermis, and DWAT in (**A**), (n=5-6). **E.-H.** Representative views with insets and quantification of B-CHP staining on P44 control and 10 days Adipo-β-cat^istab^; 10 days reversal in upper-dermis, lower-dermis, DWAT in (**E**) (n=4-5) **I., J., M., N.** Collagen was stained with picrosirius red and imaged with polarized light microscopy (red). Representative images and location of insets (white box) of genotypes in (**A., E.**). **J., N.** Insets from (**I, M**), High Density Matrix (HDM) and line masks generated by TWOMBLI algorithm plugin in ImageJ/FIJI to quantify a broad range of collagen fiber characteristics (n=6-7). (**K., L., O., P.**) Quantification of collagen fiber alignment (n=5-6) and HDM (n=6-7) based on TWOMBLI. Scale bar =100μm. *P* values were calculated with unpaired, two-tailed *t*-test with Welch’s correction and a *P* value of <0.05 is considered significant.

Topography, the surface characteristics of collagen fiber, is a good indicator of fibrotic changes in the skin (37). Based on our recent studies, we focused on the lower dermis above the DWAT and is also a comparable location across genotypes (37). Principal Component Analysis (PCA) was performed on the results of The Workflow Of Matrix BioLogy Informatics (TWOMBLI) algorithm on polarized light images of Picrosirius Red stained collagen fibers. PCA analysis showed that fiber characteristics of 10 days Adipo-β-cat^istab^ mice were very different from the control mice at the tissue level (**Supplementary figure S4A**). In addition, the high-density matrix (HDM) was significantly higher and alignment (coherency) was significantly lower in Adipo-β-cat^istab^ mice (**Fig. 3I-3L**). To further verify the alignment of the fibrotic collagen in Adipo-β-cat^istab^ mice, Alignment by Fourier Transform (AFT) algorithm was applied (**Supplementary Fig. S5A**). The heat map generated from the vector map showed that the orientation of the collagen was more disorganized and the median order parameter was significantly decreased in the Adipo-β-cat^istab^ mice (**Supplementary Fig. S5B, C**). Importantly, collagen remodeling and topography of collagen were reversible after 10 days of withdrawal from Wnt activation in the mature dermal adipocytes (**Fig. 3M-P; Supplementary Fig. S5D, E**). All together, these data indicate that Wnt activation in the mature dermal adipocytes results in elevated collagen remodeling and profibrotic collagen structure in the lower dermis next to the DWAT, which is dynamic and reversible.

### Wnt signaling activated in mature dermal adipocytes causes dermal fat loss, ECM remodeling and collagen fibrotic topography via ATGL-dependent lipolysis pathway

The lipolysis pathway, which starts with hydrolysis of triglyceride (TAG) catalyzed by ATGL to release glycerol and fatty acids, is a homeostatic pathway in adipocytes for lipid catabolism and energy generation (**Fig. 4A**). In 3T3L1 cell and mouse primary intradermal adipocyte cultures, activation of Wnt signaling with LiCl_2_ or CHIR99209 leads to lipid depletion and increase in free glycerol release (**Supplementary Fig. S6)**. This effect is attenuated by the ATGL inhibitor, Atglistatin, suggesting that the lipid depletion occurs through an ATGL-dependent pathway (**Supplementary Fig. S6)**. Next, we queried the activation of the lipolysis pathway by visualizing the expression of phosphorylated-Hormone Sensitive Lipase (p-HSL) in Adipo-β-cat^istab^ mice. Compared to undetectable of p-HSL expression in the control, we found expression after 5 days of Wnt activation in DWAT layer of Adipo-β-cat^istab^ mice (**Fig. 4B, C**). To determine if the lipolysis pathway is functionally required for DWAT lipodystrophy in Adipo-β-cat^istab^ mice, we conditionally deleted *Atgl* with the tamoxifen inducible *Adiponectin*-CreER line in mature dorsal dermal adipocytes as previously described (7,31,32) (**Fig. 4D, E**). We found nearly 33% were Myc-tag-β-catenin+ adipocytes were in Adipo-β-cat^istab^; *Atgl^fl/fl^***(Supplementary Fig. S7**). After 10 days of Wnt activation in the mature dermal adipocyte, the DWAT and dermal thickness were comparable to controls, showing that *Atgl*-dependent lipolysis is required for Wnt activated lipodystrophy of the DWAT (**Fig. 4F-4H**). The size of the individual adipocytes in Adipo-β-cat^istab^; *Atgl^fl/fl^* mice is comparable to the controls, showing the adipocytes are protected from lipodystrophy (**Fig. 4I, J**). Finally, we investigated the impact of Wnt-activation-caused-DWAT lipolysis on the ECM remodeling and collagen topography. The B-CHP expression showed comparable levels of ECM remodeling in the control and 10 days of Wnt activation in Adipo-β-cat^istab^; *Atgl^fl/fl^* mice (**Fig. 4K-4N**). Meanwhile, TWOMBLI and AFT analysis of the collagen topography showed that the HDM, alignment and median order parameters in the 10 days Adipo-β-cat^istab^; *Atgl^fl/fl^* mice were all comparable to the control (**Fig. 4O-Q**, **Supplementary Fig. S8**).

**Figure 4.**
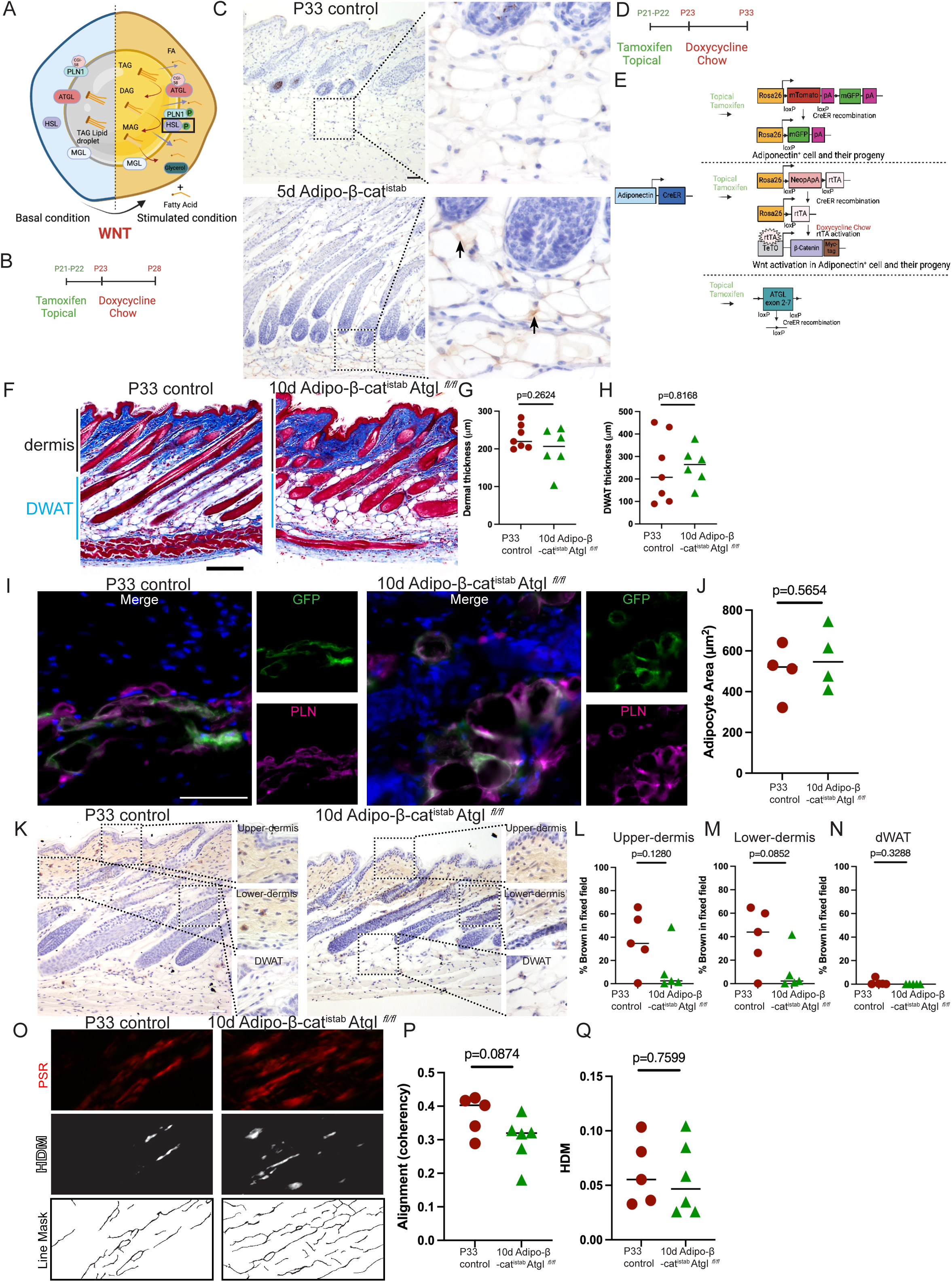
Adipo-β-cat^istab^ leads to lipodystrophy and fibrotic skin via ATGL-dependent lipolysis pathway. **A.** Schematic of lipolysis pathway. **B.** Treatment timeline of Adipo-β-cat^istab^ mice. **C.** Representative images with insets of immunohistochemistry staining of phosphorylated-HSL (brown) in control (P28) and 5 days Adipo-β-cat^istab^. Scale bar=100μm. **D.**, **E.** Treatment timeline and transgenes in the Adipo-β-cat^istab^; *Atgl^fl/fl^* mouse model. **F.** Masson’s trichrome staining of mouse dorsal skin for P33 controls CreER negative; *Atgl^fl/fl^*, 10 days Adipo-β-cat^istab^; *Atgl^fl/fl^*. Bar=200μm, black and blue bars are the dermal and DWAT thickness. **G.**, **H.** Quantification of dorsal dermal thickness and DWAT thickness in (**F**) (n=6-7). **I., J.** GFP and Perilipin 1 immunofluorescence staining and quantification of adipocyte area of GFP+ lipid droplets in skin sections of genotypes in (**F**) (n=4). DAPI=blue, GFP=green, PLN1=purple, scale bar=100μm. **K.-N.** B-CHP staining with representative views with regions of interest (as shown in Fig.3A) and quantitation of collagen remodeling staining with B-CHP (brown) in a fixed area on skin sections of genotypes in (**F**). Scale bar=200μm, (n=6). **O.** Polarized light images of dorsal dermal collagen stained with picrosirius red (PSR). High Density Matrix (HDM) and line masks images generated by FIJI on skin sections of genotypes in (**F**). **P., Q.,** Quantification of collagen fiber alignment and HDM based on TWOMBLI algorithm (n=5-6). *P* values were calculated with unpaired, two-tailed *t*-test with Welch’s correction and a *P* value of <0.05 is considered significant.

Altogether, the activation Wnt signaling in the mature dermal adipocytes results in lipodystrophy, ECM remodeling and the formation of profibrotic collagen, a process facilitated by ATGL-dependent lipolysis pathway (**Fig.5**).

**Figure 5:**
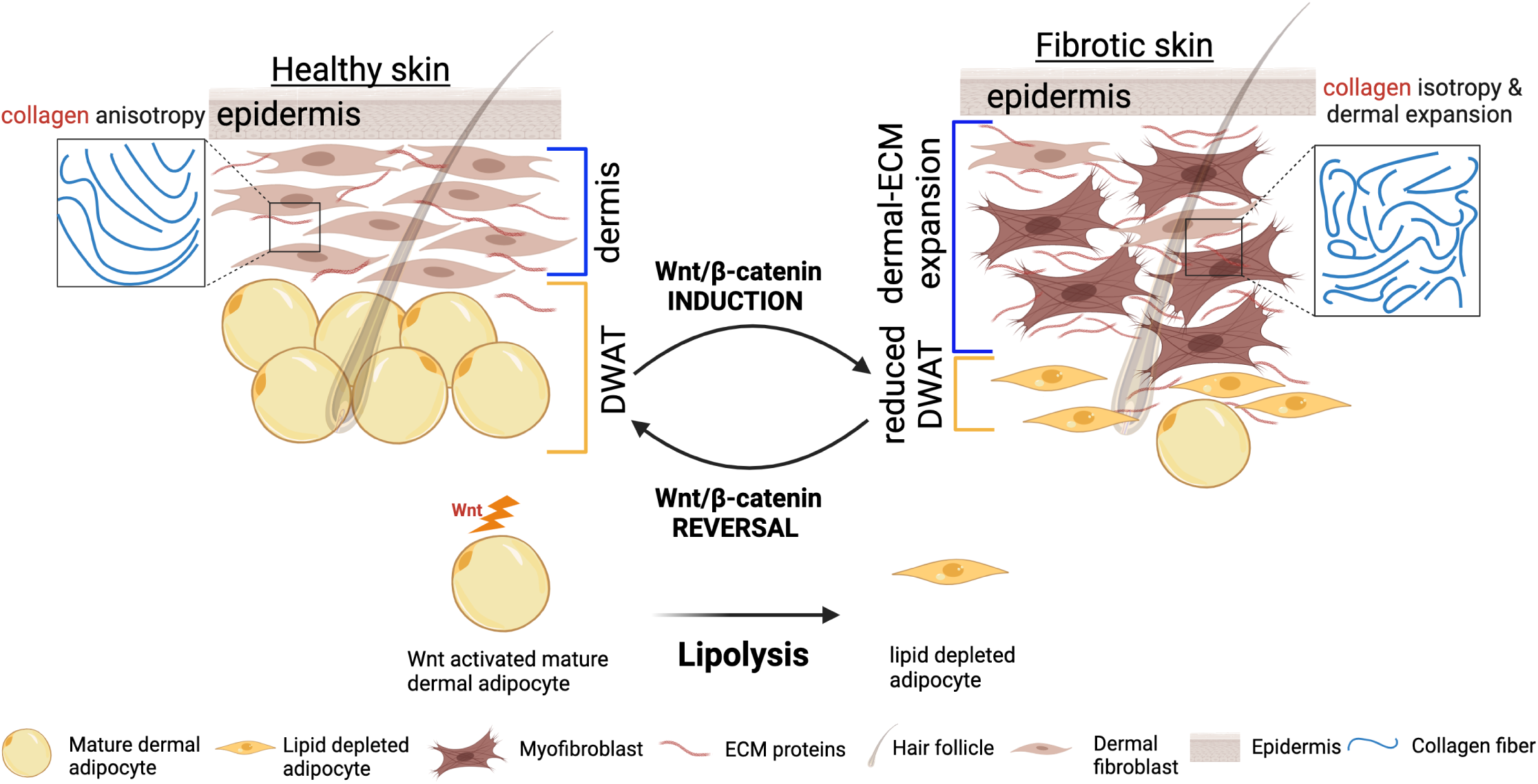
Summary model showing lipodystrophy and ECM remodeling in fibrosis are dependent on sustained Wnt activation in dermal adipocytes via ATGL-mediated lipolysis pathway.

## Discussion

By using an inducible and reversible model of Wnt activation in mouse mature dermal adipocytes, we demonstrated that lipodystrophy and associated ECM fibrotic remodeling is dependent on sustained Wnt activation in dermal adipocytes. We determined that ATGL-dependent lipolysis in the mature dermal adipocytes is required for both Wnt induced lipodystrophy and the associated ECM remodeling.

By combining lineage tracing with the inducible and reversible model of Wnt activation, we were able to determine which aspects of fibrosis and lipodystrophy are reversible in mouse skin. First, we found that Wnt induced lipodystrophy in mature dermal adipocytes is a reversible state. We found lineage marked GFP+ mature dermal adipocytes can recover in size within 10 days after withdrawal from Wnt activation. Recovery of lineage labeled dermal adipocyte cell size and DWAT area at the tissue level suggests refilling of the lipid-depleted adipocytes. The mechanism of how the adipocytes replenish the lipid content after withdrawal from Wnt activation is unclear. Either reuptake of fatty acids or use of the de novo lipogenesis pathway to accumulate lipid content could allow adipocytes to store fatty acids after Wnt activation is abrogated (38). Future studies could focus on physiological pathways that facilitate recovery of lipodystrophic adipocytes. Second, our inducible and reversible Wnt activation in dermal adipocytes revealed that profibrotic ECM remodeling in the DWAT and dermal fibroblast layer is also a reversible state. Our model demonstrates that Wnt-activation in a subset of mature dermal adipocytes not only has a cell autonomous effect, but also impacts neighboring cells and dermal layer. This interpretation is supported by the evidence of increase in cell proliferation, ECM accumulation and collagen remodeling in the DWAT and dermal layers in Adipo-β-cat^istab^ dorsal skin. It is possible that the free fatty acid and glycerol released by Wnt-induced lipodystrophy can activate the quiescent dermal fibroblasts into profibrotic fibroblasts. Studies have shown that perturbation of fatty acid oxidation to glycolysis or expression of CD36, fatty acid transporter, leads to accumulation of ECM in fibroblasts (39). Thus, our inducible and reversible model of Wnt activation in dermal adipocytes highlights the importance of lipid depletion event in the DWAT layer as a new player in promoting skin fibrosis.

The source of the profibrotic cell that mediates the ECM remodeling and accumulation in most fibrosis models and our model is unclear. There is controversy in the field about the transition of mature adipocytes to activated fibroblasts at the functional level in acute and chronic fibrosis models (40–44). In the standard bleomycin fibrosis model, a small number of non-inducible *Adiponectin*-Cre lineage labeled adipocyte cells appeared in the dermal layer with fibroblast morphology, suggesting trans-differentiation (40). Recently it was shown adipocytes have the ability to dedifferentiate during wound healing (45). Adiponectin, a signature gene of the mature adipocytes, decreases in expression during adipocyte de-differentiation (46). Adiponectin receptor signaling can downregulate collagen and α-smooth muscle actin mRNA expression in dermal fibroblasts *in vitro* (47). Our data supports a model where Wnt activation leads to lipodystrophy in dermal adipocytes that might allow them to de-differentiate to further contribute to the profibrotic process. Future research will take advantage of single nuclei RNA seq analysis to track the cell trajectory of GFP+ lineage marked cells and other cell types in the whole skin of Adipo-β-cat^istab^ mice.

The mechanisms of DWAT lipodystrophy and its impact on ECM in skin are emerging from functional studies of ATGL-lipolysis pathway. In the short-term bleomycin injection skin fibrosis model (Adipo*; Atgl^fl/fl^*) or with pharmacological inhibition with Atglistatin on wild type dorsal skin, leads to dermal expansion (48,49). In both these models, bleomycin exposure and Atglistatin treatment is on the whole skin thereby affecting multiple cell types. Here, we restricted Wnt activation and *Atgl* deletion to mature dermal adipocytes (Adipo-β-cat^istab^ *Atgl^fl/fl^*) and rescued the lipodystrophy and ECM accumulation. Our data suggests that lipolysis from dermal adipocytes are required for ECM accumulation in the DWAT and dermal layers. Our tissue restricted model will allow us to further dissect the molecular mediators that promote ECM accumulation and show that lipolysis pathway could be a new therapeutic target for modulating ECM remodeling and adipocyte lipid handling.

The mechanism of how Wnt signaling activation in dermal adipocytes activates ATGL-dependent lipolysis is unclear. It could be mediated by other Wnt effectors of lipodystrophy such as Dipeptidyl peptidase-4 (DPP4) and its substrates (18) or physiological processes such as Reactive Oxygen Species (ROS) (50). Further investigation is necessary to determine the role and mechanism of these factors in activation of lipolysis axis in our model. Clinically, recovery from the lipodystrophy during skin fibrosis could possibly bring back the functions of the DWAT layer in immune response, hair follicle cycle regulation, thermoregulation and vascularization and restore ECM homeostasis (7,10,11).

## Acknowledgements

We thank all the past and present members of the Atit, and Horsley labs for their input and feedback on this project. We thank Dr. Valerie Horsley for critical reading of the manuscript. We thank Rachel Kim, Emilia Sanz-Rios for quantification of Trichrome staining, and Dr. Maria Fernanda Fiorni for technical support on mRNA extraction of whole skin. We thank Dr. Anna Jussila for tissue culture experiments and developing some of the protocols. We thank the bio shared instrumental facility at Case Western Reserve University. This project was funded by NIH-AR076938 (Radhika Atit and Valerie Horsley) and National Scleroderma Foundation (Qiannan Ma, and Suneeti Madhavan).

## Conflicts of Interest to declare

None

## Data Sharing

Data will be made available after publication and request.

## Author Contributions

All authors were involved in drafting the article or revising it critically for important intellectual content, and all authors approved the final version to be published. Dr. Atit had full access to all of the data in the study and takes responsibility for the integrity of the data and the accuracy of the data analysis.

**Study conception and design**: Atit and Ma

**Acquisition of data**: Ma, Segal, Reynolds, Gregory, Wyetzner, Madhavan

**Analysis and Interpretation of data**: Atit, Ma, Segal, Wyetzner, Madhavan

---

**Supplementary figure 1.** Specificity of *R26mTmG* recombination Adipo-β-cat^istab^ mice. **A.** GFP immunofluorescence staining of gonadal (Lane 1 left), Inguinal (Lane 1 right) and Subcutaneous (Lane 2 left) in *Adiponectin*CreER/+; R26mTmG/+ mouse. DAPI=Blue, GFP=Green, mTomato=red, scale bar = 100μm **B.** Quantification of percentage of GFP positive adipocytes vs mTomato positive adipocytes in AdiponectinCreER/+; R26mTmG/+ mouse DWAT, Gonadal, Inguinal and Subcutaneous (n=3-5). **C-D.** GFP and Perilipin 1 immunofluorescence staining in control (P33) with dermis and between hair follicles. * non-specific staining. *P* values were calculated with unpaired, two-tailed *t*-test with Welch’s correction and a *P* value of <0.05 is considered significant.

**Supplementary figure 2.** Wnt signaling pathway is efficiently activated in mature dermal adipocytes in the 5 days Adipo-β-cat^istab^ mice. **A.** Immunohistochemistry staining of Myc-tag on mouse dorsal skin in both dermis (up) and DWAT (bottom) on P28 control and 5d Adipo-β-cat^istab^; scale bar =100μm **B.** Schematic of immunohistochemistry staining and detection of Myc-tag **C.** Quantification of Myc-tag positive adipocytes vs control (n=4-7) **D.** Quantification of Axin2 mRNA level by qPCR in P28 controls and 5d Adipo-β-cat^istab^. *P* values were calculated with unpaired, two-tailed *t*-test with Welch’s correction and a *P* value of <0.05 is considered significant.

**Supplementary figure 3.** Cell survival and proliferation in Adipo-β-cat^istab^ mice. **A.** TUNEL staining for P28 control, 5 days Adipo-β-cat^istab^ and P42 hair follicle as positive control. DAPI=blue, TUNEL=red, PLN1=green, scale bar= 100μm. **B.** TUNEL staining for P33 control and 10 days Adipo-β-cat^istab^. DAPI=blue, TUNEL=red, PLN1=green, scale bar= 100μm. **C.** Ki67 and GFP immunostaining 10 days Adipo-β-cat^istab^ between hair follicles with the white arrows indicate the Ki67+ signal, scale bar= 100μm. **D.** Ki67 and GFP immunostaining 10 days Adipo-β-cat^istab^ in the dermis with the white arrows indicate the epidermis Ki67+ signal and yellow arrows indicate dermis Ki67+ signal, scale bar=100μm.

**Supplementary figure 4.** Adipo-β-cat^istab^ leads to changes in some parameters of fibrotic topography of collagen. **A.** Principal component analysis (PCA) plots to TWOMBLI metrics of P33 control and 10 days Adipo-β-cat^istab^. Each point represents one ROI and 12 images were taken for each animal (n=5-7). **B.-D.** Quantification of lacunarity, number of endpoints, hyphal growth unit (HGU) on P33 control and 10 days Adipo-β-cat^istab^, (n=5-7). **E.** Principal component analysis (PCA) plots to TWOMBLI metrics of P43 control and 10 days Adipo-β-cat^istab^; 10 days reversal. Each point represents one ROI and 12 images were taken for each animal (n=6). **F.-H.** Quantification of lacunarity, number of endpoints, hyphal growth unit (HGU) on P43 control and 10 days Adipo-β-cat^istab^; 10 days reversal, (n=6). *P* values were calculated with unpaired, two-tailed *t*-test with Welch’s correction and a *P* value of <0.05 is considered significant.

**Supplementary figure 5.** Adipo-β-cat^istab^ leads to less alignment of collagen shown by Alignment of Fourier Transform (AFT). **A.** Schematic of workflow and output of AFT based on picrosirius red staining of collagen. **B.** Vector maps (top) of P33 control and 10 days Adipo-β-cat^istab^ generated from 16-bit images of 40x polarized light image cropped ROI. Yellow bar stands for the orientation of the collagen. Heatmaps of P33 control and 10 days Adipo-β-cat^istab^ based on the vector maps. **C.** Quantification of median order parameter on P33 control and 10 days Adipo-β-cat^istab^ (n=6). **D.** Vector maps (top) of P43 control and 10 days Adipo-β-cat^istab^; 10 days reversal generated from 16-bit images of 40x polarized light image cropped ROI. Yellow bar stands for the orientation of the collagen. Heatmaps of P43 control and 10 days Adipo-β-cat^istab^ 10 days reversal based on the vector maps. **E.** Quantification of median order parameter on P43 control and 10 days Adipo-β-cat^istab^; 10 days reversal, (n=6-7). *P* values were calculated with unpaired, two-tailed *t*-test with Welch’s correction and a *P* value of <0.05 is considered significant.

**Supplementary Fig. 6:** Wnt activation leads to lipid depletion in dermal adipocytes via ATGL *in vitro* **A.** 3T3-L1 adipocytes treated with differentiation media, or treated with differentiation media followed by 7μM Wnt agonist, LiCl_2_, or treated with differentiation media, LiCl_2_, and 40μM ATGL inhibitor, Atglistatin; scale bar=200μm. **B.** Primary intradermal adipocyte progenitor cells untreated, or treated with differentiation media, or treated with differentiation media followed by 7μM Wnt agonist, CHIR99021, or treated with differentiation media, CHIR, and 40μM ATGL inhibitor, Atglistatin, or treated with differentiation media, followed by 40μM Atglistatin; scale bar=200μm; average Oil Red O area (2 fields per well, 4 biological replicates, and 2-4 technical replicates per sample per treatment) and free glycerol quantification (2 technical replicates and 4 biological replicates/sample). *P* values were calculated with unpaired, two-tailed *t*-test with Welch’s correction and a *P* value of <0.05 is considered significant.

**Supplementary figure 7.** Wnt signaling pathway is efficiently activated in mature dermal adipocytes in Adipo-β-cat^istab^; *Atgl ^fl/fl^* mice. **A.** Immunohistochemistry staining of Myc-tag on mouse dorsal skin DWAT on P33 control CreER negative; *Atgl^fl/fl^*and 10d Adipo-β-cat^istab^ *Atgl ^fl/fl^*. Scale bar = 100μm **B.** Quantification of Myc-tag positive adipocytes vs control (n=4-5). *P* values were calculated with unpaired, two-tailed *t*-test with Welch’s correction and a *P* value of <0.05 is considered significant.

**Supplementary figure 8.** Collagen topography changes is comparable to controls in Wnt activated-dermal adipocyte conditional *Atgl* mutants (Adipo-β-cat^istab^; *Atgl^fl/fl^*) **A.** Principal component analysis (PCA) plots to TWOMBLI metrics of control (P33) (red), 10 days Adipo-β-cat^istab^; *Atgl^fl/fl^* (blue) Each point represents one ROI and 12 images were taken for each animal (n=5-6). **B-D.** Quantification of lacunarity, number of endpoints, hyphal growth unit (HGU) on genotypes in (A) (n=5-6). **E.** Vector maps (top) from 16 bit images of 40x polarized light image cropped ROI of genotypes in (A). Yellow bar stands for the orientation of the collagen. Heatmaps are based on the vector maps. **F.** Quantification of median order parameter for alignment of collagen fibers (n=5-6). *P* values were calculated with unpaired, two-tailed *t*-test with Welch’s correction and a *P* value of <0.05 is considered significant.

