## supplementary figures for "Wnt activation in mature dermal adipocytes leads to lipodystrophy and skin fibrosis via ATGL-dependent lipolysis"

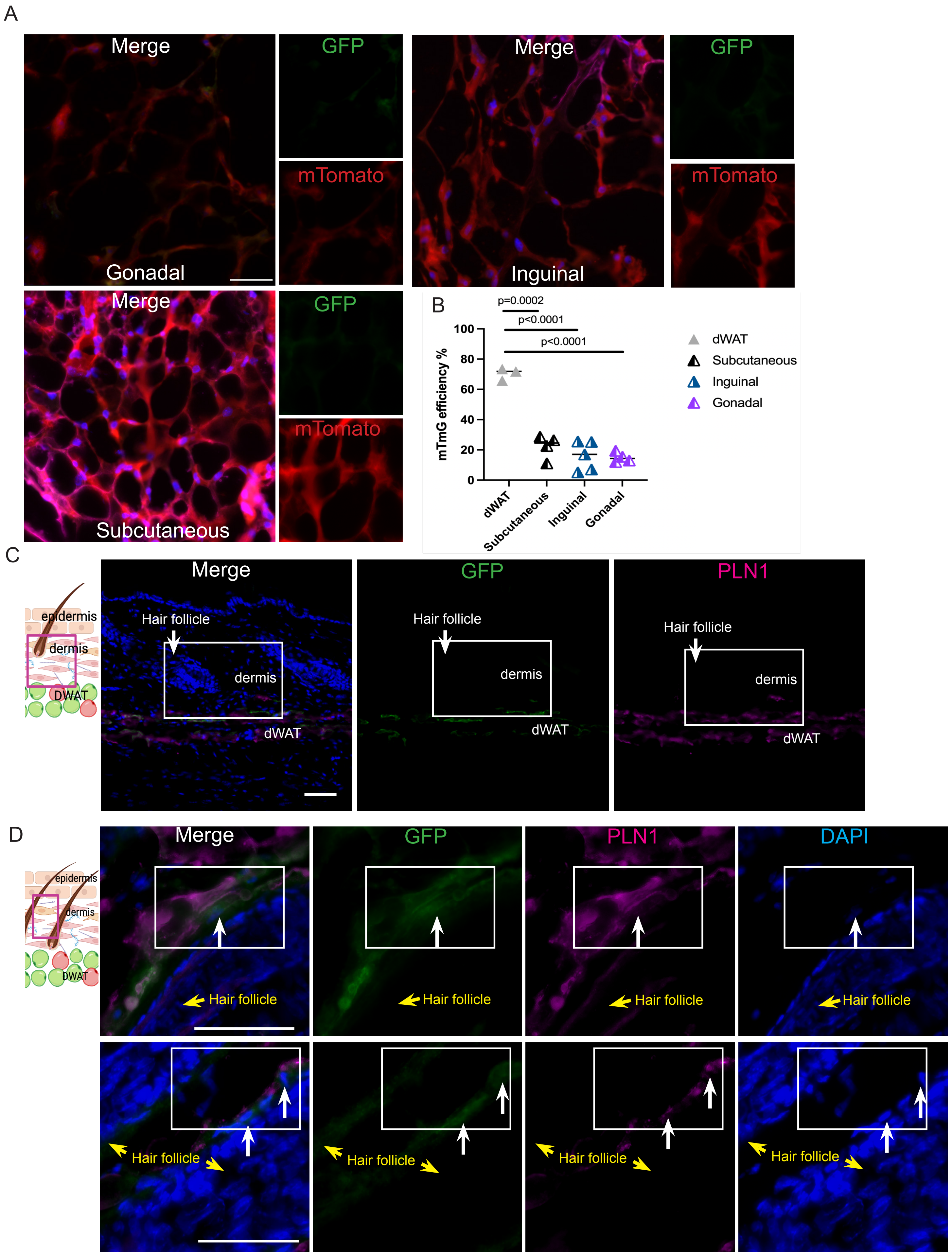


Ma et al.,
Supplementary Figure 1

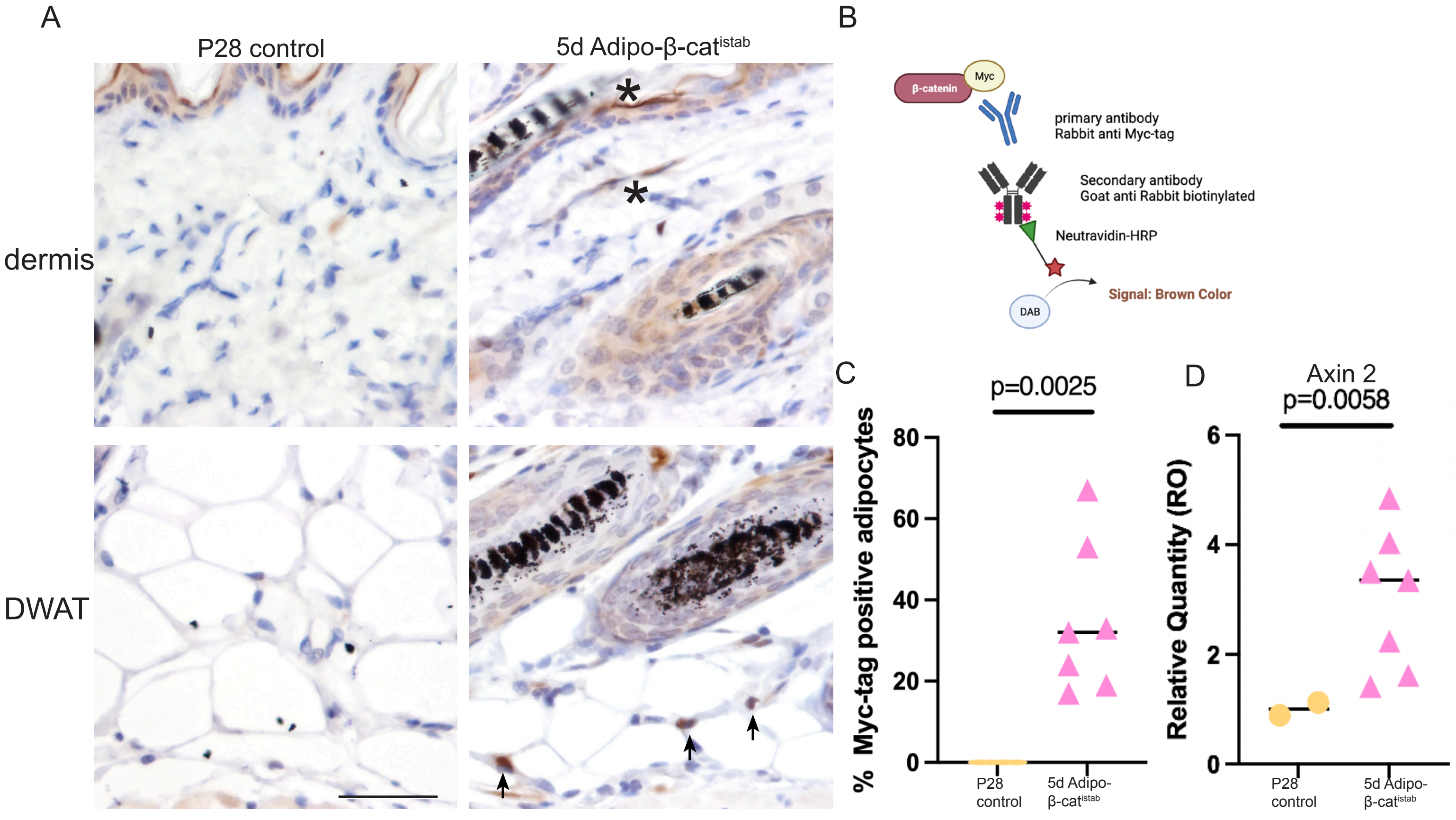


**Ma et al.,**
**Supplementary Figure 2**

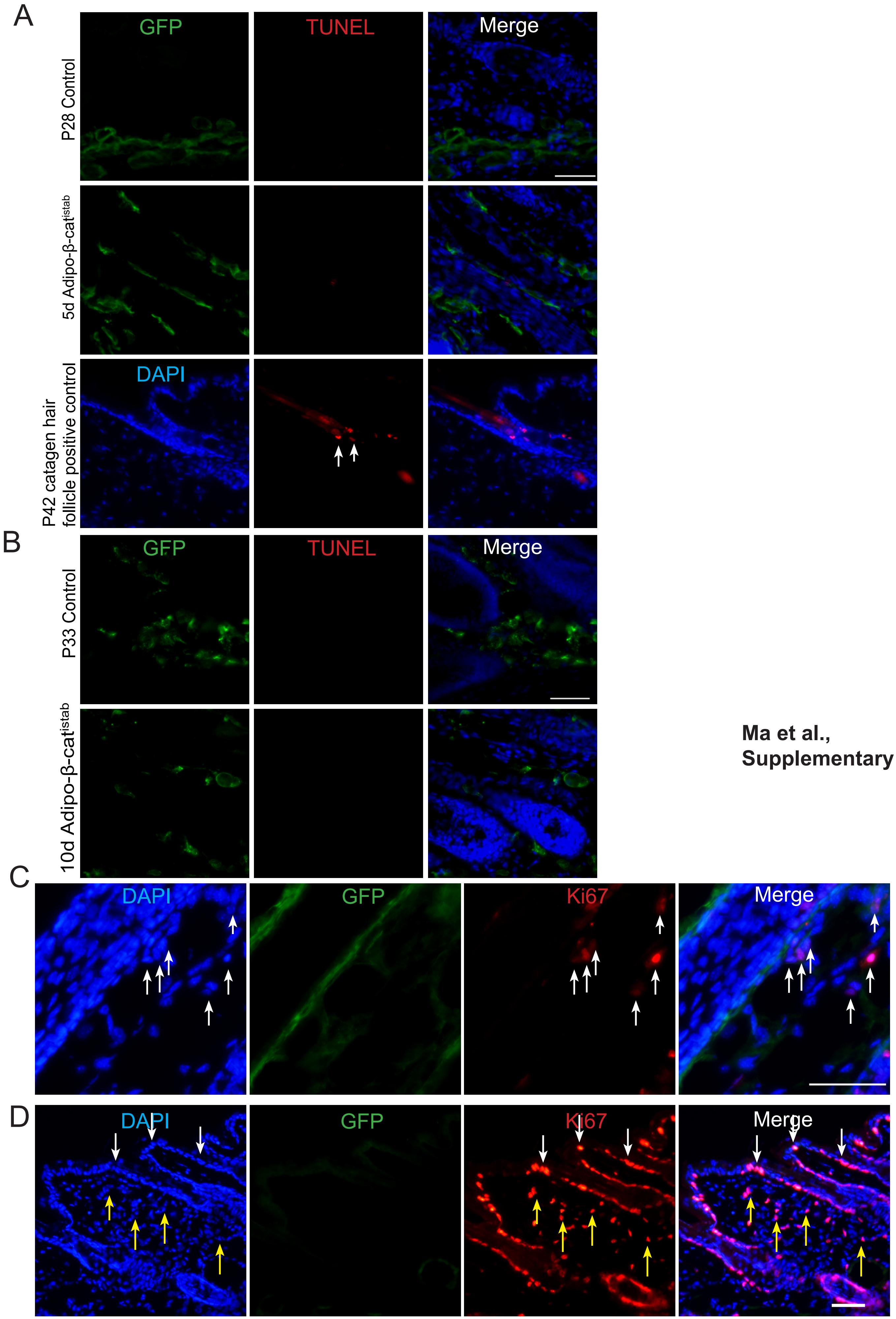


**Ma et al.,**
**Supplementary Figure 3**

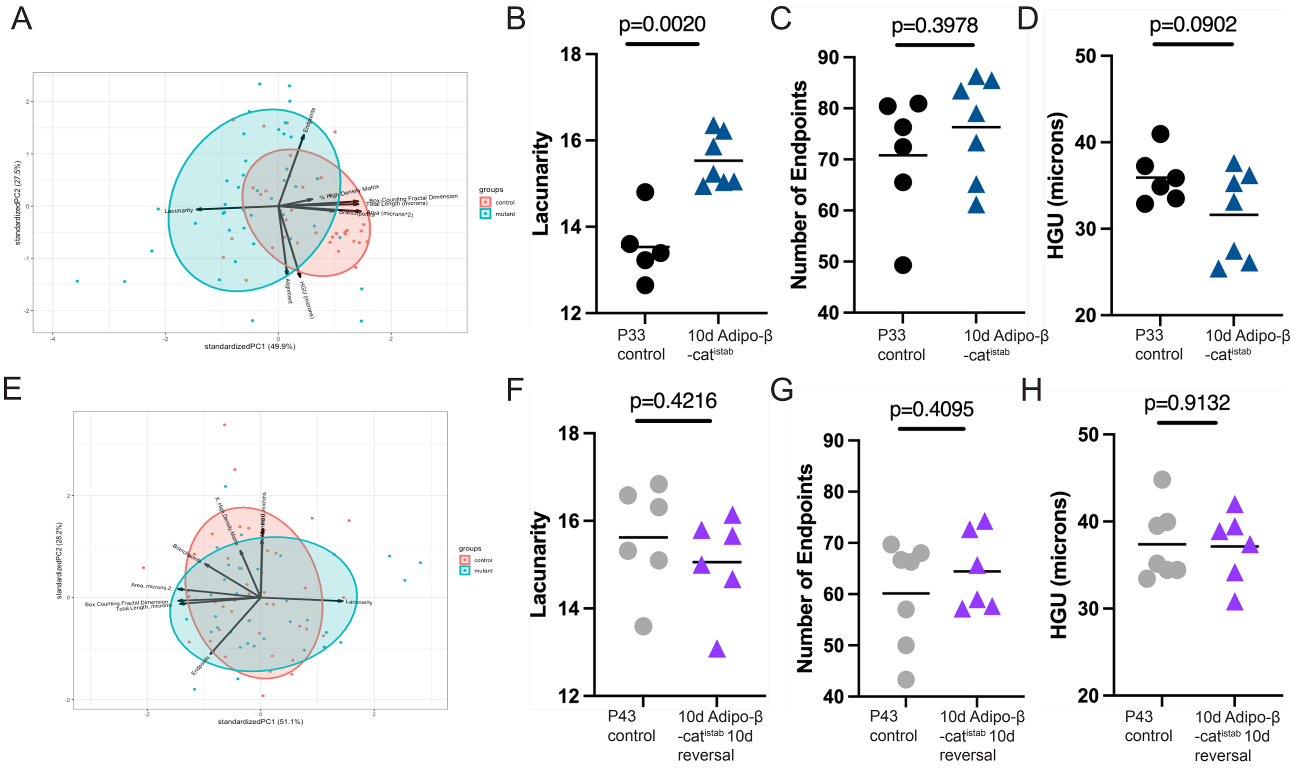


**Ma et al., Supplementary Figure 4**

**Ma et al.,**
**Supplementary Figure 5**

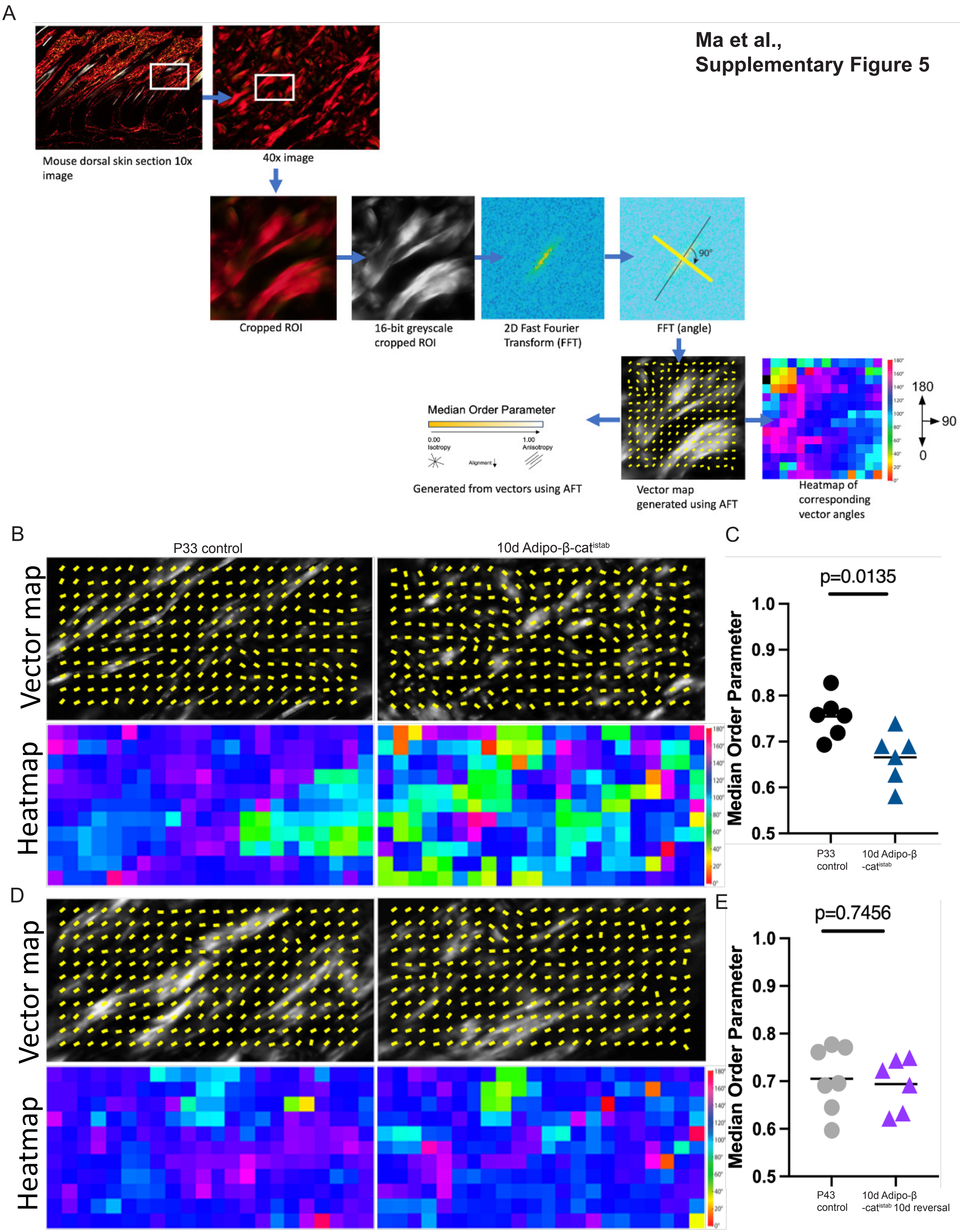

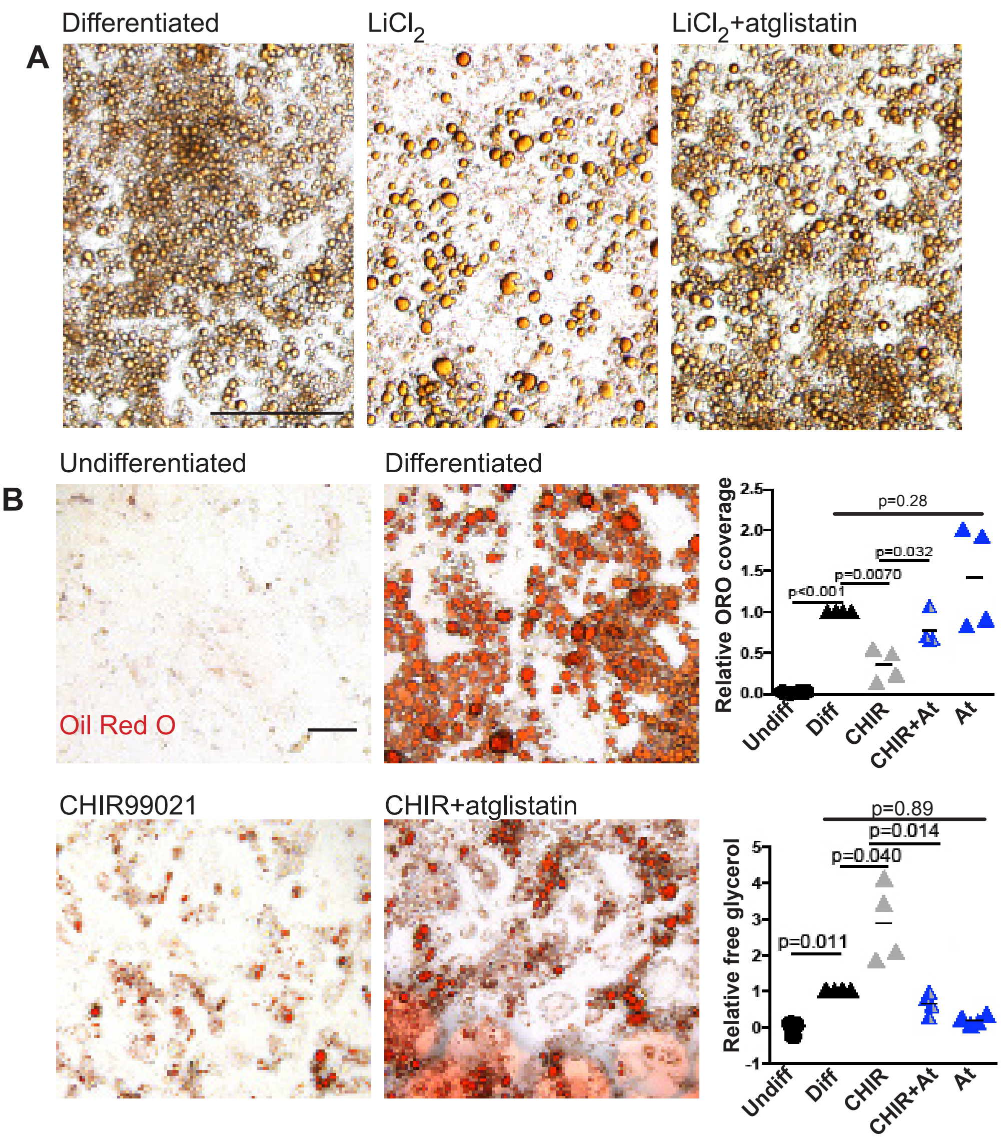


Ma et al.,
Supplementary Figure 6

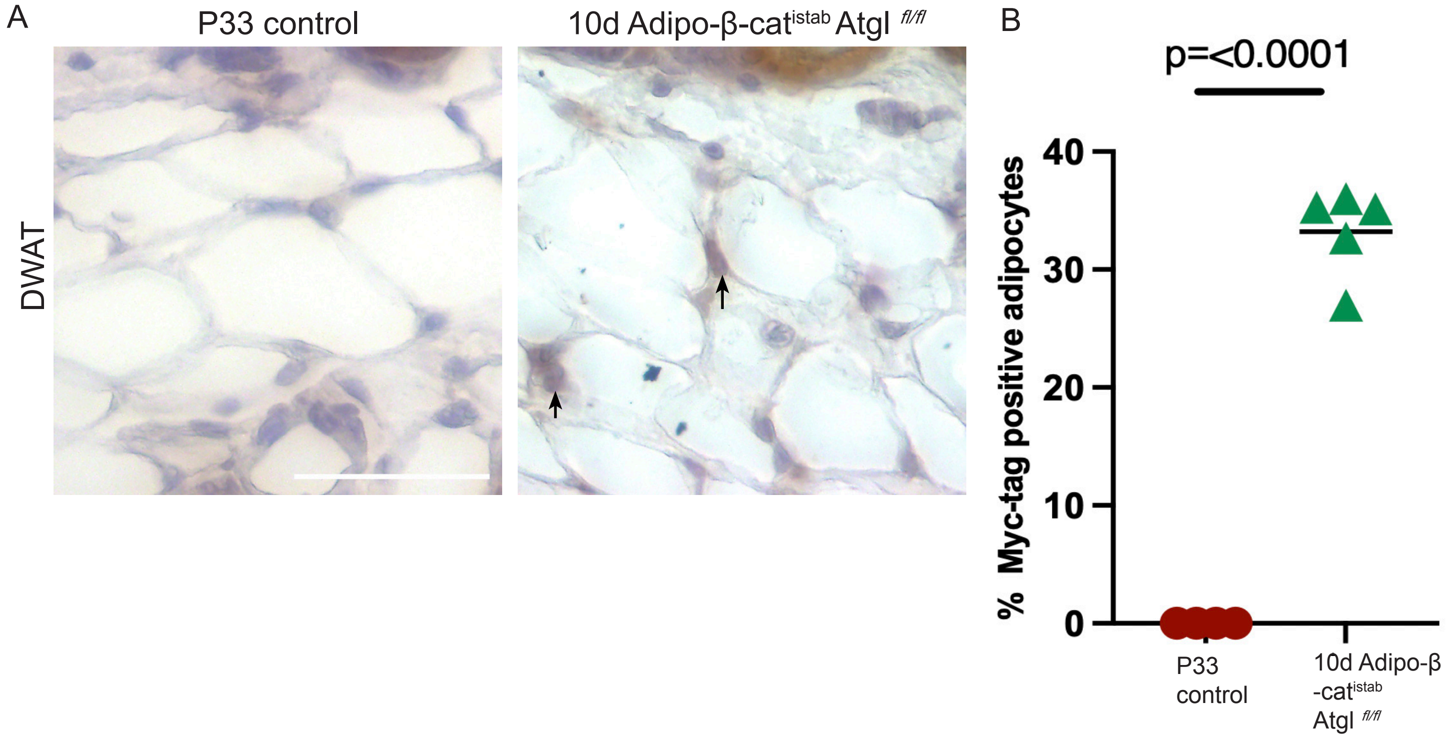


**Ma et al.,**
**Supplementary Figure 7**

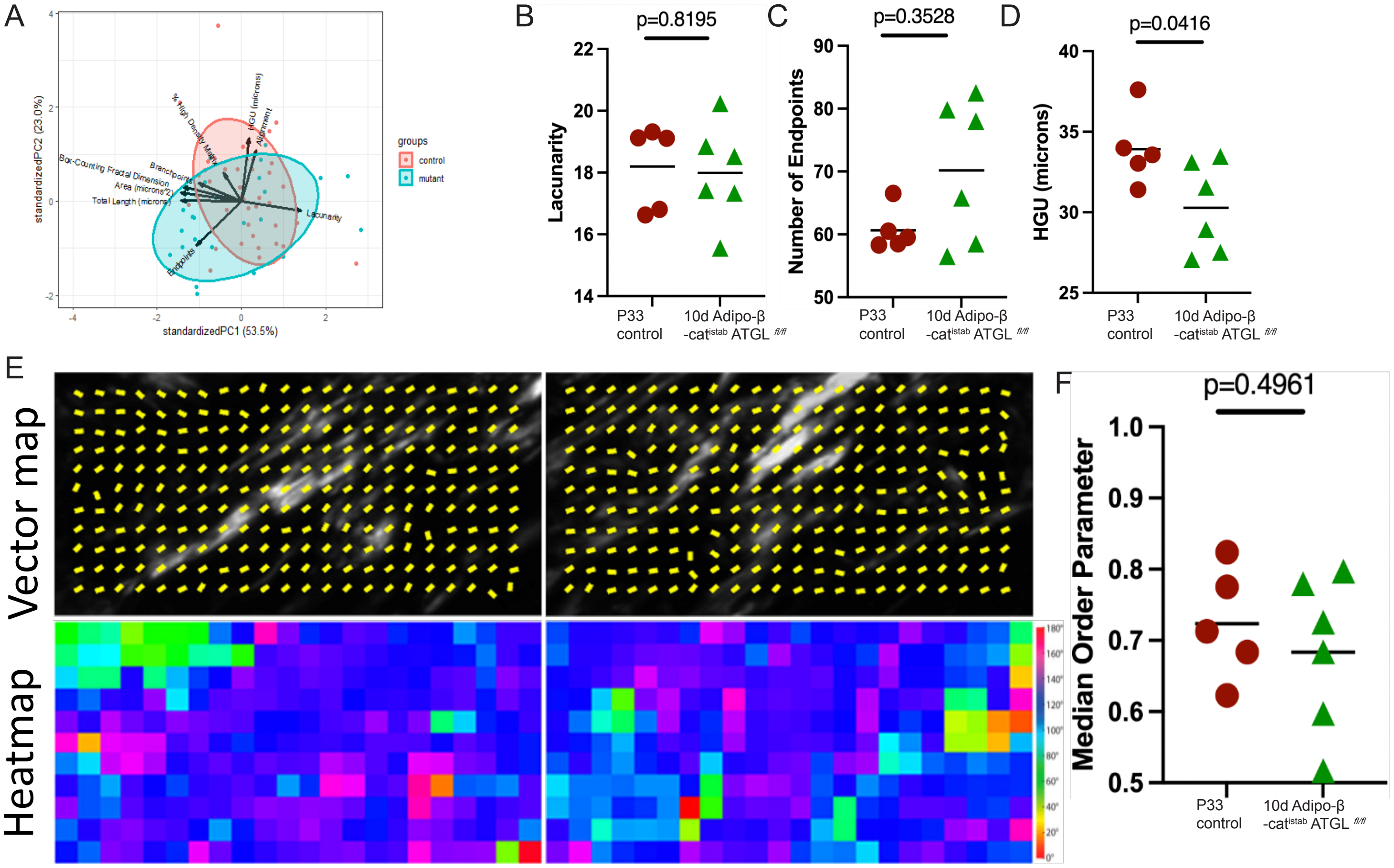


**Ma et al.,**
**Supplementary Figure 8**
