## Supplementary methods for "Wnt activation in mature dermal adipocytes leads to lipodystrophy and skin fibrosis via ATGL-dependent lipolysis"

**Histological staining and morphometrics**

Tissue was fixed in 10% buffered formalin (Fisher Scientific SF100-4) for 1 hour at 4ºC, embedded in paraffin, and sectioned at 7µm. Paraffin sections were stained with Masson's trichrome according to standard protocols. Paraffin sections were also stained with 1.6% w/v Picrosirius red (26357-02; Electron Microscopy Science) for 20 minutes then rinsed for 5 minutes in acidified water (1% acetic acid in tap water). Polarized light microscopy allows for better viewing of collagen fibers due to its birefringent nature. The Olympus BX microscope and Cell Sens entry software were used to take brightfield images. Fiji/ImageJ (National Institute of Health, Bethesda, MD) was used in phenotypic data quantification analysis for dermal thickness, dermal white adipose tissue thickness, cell counting, and individual adipocyte area measurements. Quantification of skin compartment thickness represents the average of three different locations (left, middle and right) in 3-6 sections/mouse.

**Immunohistochemistry and immunofluorescence**

Paraffin tissue was fixed in 10% formalin for 1 hour at 4ºC, embedded in paraffin, and sectioned at 7µm. Paraffin sections were deparaffinized by successive ethanol solutions of 99%, 96%,90%, 80%, 70%, and 50%, then washed in 1x PBS. Then slides underwent heat-based antigen retrieval in a citrate buffer (10 mM Tri-Sodium Citrate dihydrate, pH=6) at 90-94ºC for 15 minutes in a water bath. Slides were then cooled for 50 minutes to room temperature and blocked with 10% goat serum (Thermo Fisher Scientific 16210064), 1% BSA (BP1600100; Fisher Scientific), and 0.05% Tween 20 (BioWorld 42030016-1) for 85 minutes at room temperature in a humid chamber. Tissues were incubated with proper primary antibodies diluted to a certain ratio with blocking buffer in a humid chamber at 4ºC overnight. The following primary antibodies were used for bright field immunohistochemistry and immunofluorescence as described previously (1–3). Rabbit anti-Myc Tag (ab9106, 1:500; Abcam), Rabbit anti phospho-HSL (Ser 565) (4137, 1:1600; Cell Signaling), Rabbit anti-alpha SMA (ab124964, 1:1000; Abcam). For Myc-tag and p-HSL, after antigen retrieval by citric buffer, 0.3% $H_2O_2$ for 10 minutes, 5 minutes TBS wash and 85 minutes blocking buffer (1% goat serum+1% BSA+ 0.1% Tween 20 in TBS) incubation, Goat anti-Rabbit Biotinylated secondary antibody (1:250; Vector Labs, San Francisco, CA) was used for 30 minutes at room temperature. After two 5 minutes washes in TBS, 4µg/mL neutravidin-HRP (31030; Thermo Scientific) diluted in a blocking buffer for 30 minutes was applied followed by DAB development (Myc-tag is 6.5 minutes, p-HSL is 10 minutes). Nuclei were counterstained with hematoxylin before mounting in Permount Mounting Medium (SP15-100; Fisher Chemical) or counterstained with DAPI (D9542,1:2000; Sigma-Aldrich) before mounting in Fluoroshield (Sigma-Aldrich). Positive and negative controls were used for specificity detection of the antibodies.

Flash-frozen dorsal skin is embedded in OCT (Tissue-Tek) and sectioned at 14µm at -20ºC. Cryo-sectioned samples were fixed by 4% PFA for 15 minutes at room temperature and washed in PBS twice. Slides were blocked with 10% Donkey serum (50413115; Fisher Scientific) and 0.25% Triton X-100 (BP151100; Fisher Scientific) for 30 minutes at room temperature in a humid chamber. Tissues were incubated with proper primary antibodies diluted in block buffer to a certain ratio in a humid chamber at 4ºC overnight. The following primary antibodies were used for immunofluorescence as described previously (3,4) : Chicken anti GFP (ab13970, 1:1000; Abcam), Rabbit anti Perilipin (ab3526, 1:500; Abcam), Rabbit anti Ki67 (ab15580, 1:500; Abcam). After two 5-minute PBS washes, Donkey anti Rabbit Alexa 647 (A31573,1:500; Thermo Fisher Scientific), Donkey anti Chicken Alexa 488 (111545003, 1:500; Jackson ImmunoResearch), and Donkey anti Rabbit Alexa 488 (A32790, 1:500; Thermo Scientific) secondary antibodies were applied at room temperature for 30 minutes. TUNEL staining was performed on Cryosections by TUNEL TMR red kit (11767291910, 5% TUNEL enzyme; Roche Diagnostic Crop). Nuclei were counterstained with DAPI and then mounted in Fluoroshield. Positive and negative controls were used for specificity detection of the antibodies and kit.

**B-CHP staining**

Paraffin sections were deparaffinized by successive ethanol solutions then washed in PBS. Slides were then blocked with 5% goat serum in PBS at room temperature for 20 minutes. Apply 3-4 drops per slide of biotin block (SP 2001; Vector Laboratories) and incubate the slides in the humid chamber for 15 minutes at room temperature. Stain with a biotin-conjugated collagen hybridizing peptide (B-CHP, BIO300, 3Helix) at a concentration of 2μM in block buffer overnight. 4μg/mL neutravidin-HRP diluted in a blocking buffer was applied for 30 minutes followed by 4 minutes of DAB (SK-4100; Vector Laboratories) development.

**Imaging**

Masson's Trichrome, Myc-Tag, p-HSL, and Picrosirius red images were taken at room temperature by an Olympus BX60 microscope with a digital camera (DP70, Olympus). Masson's trichrome staining images were taken at 4x objective (Olympus UPlanFI 4x/0.13) by Cell Sens Entry software (Version 1.5, Olympus Corporation 2011). Myc-Tag and p-HSL images were taken at 10x and 40x objectives. Picrosirius red images were taken at 40x under polarized light produced by microscope U-ANT (U-P115, Olympus) and U-POT (U-P110, Olympus). Exposure between controls and mutants was the same. Immunofluorescence images were taken by an inverted wide-field Leica Dmi8 microscope at 20x (506243 Germany) and 40x (506243 Germany) magnification. Images were analyzed in Fiji/ImageJ and processed by Adobe Photoshop and Adobe InDesign.

**Morphometric and image analysis pipelines:**

*Adipocyte area measurement*
20x immunofluorescence images of GFP and PLN1 double staining were used for individual adipocyte area measurement. Approximately 30 adipocytes per mouse of control and GFP+ PLN1+ mutant were measured from non-overlapping fields and analyzed using the segmentation editor tool

in Fiji/ImageJ. The areas of adipocytes were binned into a histogram using GraphPad Prism (GraphPad Software, San Diego, CA).

*Collagen amount measurement*
10x Masson's Trichrome stained images were used for total collagen analysis of the DWAT. The blue which stains for collagen in the Masson's Trichrome staining was threshold by ImageJ with the HSB parameters (Hsu: 135-195; Saturation: 25-255; Brightness: 20-255) (5). The white color in the mask image after HSB threshold stands for the blue stained collagen. For each animal, 4-6 images were used and for each image, three 40 pixels x 30 pixels ROI were taken at the bottom of the DWAT and the middle of the DWAT. The area of the white color in the ROI was collagen amount.

*Cell proliferation analysis*
For quantification of Ki67 positive cells in epidermis and dermis, 10x immunofluorescence images of Ki67 and GFP double staining were used. Measurements were made in rectangular ROIs (170x144px) in the left, middle and right of the dermis or epidermis of each image. The number of nuclei with overlapping DAPI and Ki67 nuclei were counted with Fiji's cell counter tool. The ratio of overlapping Ki67 and DAPI nuclei to all DAPI stained nuclei was taken. For each animal (controls and mutants), 3 images were used from left, middle and right of the tissue section and on each image one ROI on the left, middle and right of the picture were used. For quantification of Ki67 positive cell number in DWAT, 40x immunofluorescence images of Ki67 and GFP double staining were used. Fiji cell counter was used to count GFP+ adipocyte and Ki67+ nuclei within GFP+ adipocytes. Ki67 nuclei to total GFP positive adipocytes was taken as the ratio of Ki67+ in the DWAT. 3 images were used from left, middle and right of the tissue section for each mouse (controls and mutants). The results were graphed as a scatter plot using GraphPAD Prism.

*Biotin Conjugate Collagen Hybridizing Peptide (B-CHP) Analysis*
40X magnification images were used for quantification. Images were all white-balanced in Adobe Photoshop. One 350x200px ROI was taken per 40x image. Images were deconvoluted using Fiji. All images were threshold to 1,150. Percent area in brown in the ROI was taken and averaged per animal. Each animal is represented by 9 ROIs in total, 3 ROIs per each location: upper dermis, lower dermis, and DWAT.

*Myc-tag expressing adipocytes*
40x brightfield images were analyzed for Wnt activation efficiency in mature dermal adipocytes. One image from left, middle and right of the dorsal skin was analyzed per mouse. The cell counter tool of Fiji was used to count the total number of adipocytes in the DWAT area and the number of the Myc-Tag positive adipocyte nuclei (brown) inside the fixed field. The number of Myc-Tag positive adipocytes was divided by the total number of the adipocytes in the DWAT area to get the Wnt activation efficiency in the mature dermal adipocytes and graphed by GraphPAD Prism.

40x images of immunofluorescence taken by Leica Dmi8 microscope were analyzed for tamoxifen specificity. Images of sections in the DWAT, subcutaneous, gonadal, and inguinal fat were taken after topical tamoxifen application on dorsal skin and doxycycline treatment. Images were taken in the left, middle and right side across three sections on a slide and analyzed. The number of GFP positive adipocytes and total number of adipocytes in the whole image were counted by Fiji (dimension 1360x1024). The Tamoxifen recombination efficiency is the ratio of GFP positive adipocytes to the total number of adipocytes and was graphed using GraphPAD Prism.

*TWOMBLI (The Work Flow Of Matrix BioLogy Informatics) analysis of matrix*

PSR-stained mouse dorsal skin sections were imaged using polarized light microscopy and used for TWOMBLI analysis. For each animal, 3 non-overlapping 40x images from left, middle and right of the dorsal skin were taken at the dermis immediately above the DWAT in controls (P33 or P44) and from the remodeled dermis (very close to the remaining DWAT) above the panniculus carnosus muscle layer in 2 days Tamoxifen painting (P21-P22) 10 days doxycycline chow (P23-p33) and 2 days Tamoxifen painting (P21-P22) 10 days doxycycline chow (P23-p33) 10 days reversal (P34-P44). Two fix-sized of ROIs (376x195 pix) were taken from the 40 images. The ROIs from these pictures were run through TWOMBLI in Fiji/ImageJ (6). The parameters were set as the following list: Contrast Saturation: 0.35, Minimum Line Width: 5, Maximum Line Width: 10, Minimum Curvature Window: 20, Maximum Curvature Window:70, Minimum Branch Length: 10, Maximum Display HDM: 220, Minimum Gap Diameter: 9. Line masks of the matrix network were generated by the Fiji Ridge Detection tool. According to the line mask, The ECM metrics including curvature, fractal dissension, number of endpoints, lacunarity and alignment were calculated on each ROI and get the average of the total of 6 ROIs per animal.

*AFT (Alignment by Fourier Transform) analysis of collagen alignment*

The same 40x images of PSR stained mouse dorsal skin by polarized light microscopy were used to take new ROIs for AFT. These new ROIs were then converted to 16-bit grayscale. 40x non-overlapping images were used from left, middle and right from the sections. Two fixed-size of ROIs (376x195 pix) were taken from the dermis immediately above the DWAT in controls (P33 or P44) and from the remodeled dermis (very close to the remaining DWAT) above the panniculus carnosus muscle layer in 2 days Tamoxifen painting (P21-P22) 10 days doxycycline chow (P23-p33) and 2 days Tamoxifen painting (P21-P22) 10 days doxycycline chow (P23-p33) 10 days reversal (P34-P44). The ROIs were converted into 16-bit grayscale by Fiji and input into Alignment by Fourier Transform (AFT) (7) by MATLAB (Mathworks, v2023b). Median order parameter and heatmap of AFT vector orientation were output using the following parameters: Window Size: 30 pixels, Window Overlap: 50%, Neighborhood Radius: 2x vectors. Local masking and filtering were not applied. The Median Order Parameter has a range of 0 to 1 where 0 is indicative of unorganized alignment (isotropy) and 1 is complete alignment (anisotropy). The median order parameters were averaged across 6 ROIs per animal and graphed by GraphPAD Prism.

*PCA (Principal Component Analysis)*
PCA was comprised of TWOMBLI output and the covariance between variables was checked in RStudio. Each data point is one ROI with outliers removed before analysis calculated by Microsoft Excel. The curvature metrics output by TWOMBLI were not included in the PCA analysis as the result of low covariance values. PCA biplot was generated by principal components 1 and principal component 2.

**RNA extraction and qPCR analysis**

For mature dermal adipocyte qPCR, DWAT tissue was manually cut from 5 mm biopsy punches (9033515, Premier Uni-Punch) from the mouse dorsal skin after 2 days Tamoxifen painting (P21-P22) and 5 days (P23-P28) or 10 days doxycycline chow(P23-P33). RNA was extracted from flash frozen DWAT tissue by Trizol reagent (15596026; Thermo Fisher Scientific). 4ng cDNA was used for qPCR after RT reaction as previously described (2,8) Adiponectin and Axin2 mRNA were quantified relative to Hprt using Taqman master mix (4304437, Thermo Fisher Scientific) by the probes (Mm00456425_m1, Mm00443610_m1, Mm03024075_m1; Thermo Fisher Scientific).

For the whole skin qPCR, flash frozen mouse whole dorsal skins were grounded down to a fine powder in the liquid nitrogen. 350μL of Trizol (15596026, Invitrogen) was added for each 0.5cm x 0.5cm skin sample then use the RNeasy Mini Kit (74104, Qiagen) to extract RNA. 40ng cDNA was used for qPCR after RT reaction. Adiponectin and Perilipin mRNA were quantified relative to ACTB using TaqMan master mic (4304437, Thermo Fisher Scientific) by the probes (Mm00456425_m1, Mm00558672_m1, Hs01060665_g1, Thermo Fisher Scientific.

The qPCR was performed on an Applied Quantstudios Biosystem 3 PCR System. A relative quantity of fold change of Ct values controls was expressed by the 2-ΔΔCt formula. qPCR data was presented in univariate scatter plots as previously described (9) in GraphPAD Prism.

**Cell culture:**
3T3-L1 cell lines and primary intradermal adipocyte progenitors were differentiated, stained with Oil Red O, and free glycerol assay was performed as previously described (1). 3T3-L1 adipocytes treated with differentiation media,or treated with differentiation media followed by 7μM Wnt agonist, $LiCl_2$, or treated with differentiation media, $LiCl_2$, and 40μM ATGL inhibitor, atglistatin (Sigma Aldrich, SML1075) for six days and stained with Oil Red O as previously described (1). Following differentiation, primary intradermal adipocyte progenitor cells were treated with differentiation media followed by 7μM Wnt agonist, CHIR99021, or treated with differentiation media, CHIR, and 40μM ATGL inhibitor, atglistatin, or treated with differentiation media, followed by 40μM atglistatin. Cultures were stained with Oil Red O and the average Oil Red O area (2 fields per well, 4 biological replicates,

and 2-4 technical replicates per sample per treatment) was obtained using Image J/Fiji. Free glycerol release in conditioned media was quantified using manufacture instructions (Sigma Aldrich F6428) (2 technical replicates and 4 biological replicates/sample).

**Statistical analysis**

No statistical method was performed to determine the sample size. The sample size was determined by previous studies (1). Each individual point in the figures was the average value of 3-15 technical replicates per animal according to different experiments (indicated within the methods of each assay).
